# Identification of novel neuroprotective loci modulating ischemic stroke volume in a cross between wild-derived inbred mouse strains

**DOI:** 10.1101/591669

**Authors:** Han Kyu Lee, Samuel J. Widmayer, Min-Nung Huang, David L. Aylor, Douglas A. Marchuk

**Affiliations:** Department of Molecular Genetics and Microbiology, Duke University Medical Center, Durham, North Carolina, USA; Department of Biological Sciences, North Carolina State University, Raleigh, North Carolina, USA; Division of Cardiology, Department of Medicine, Duke University Medical Center, Durham, North Carolina, USA

**Keywords:** Ischemic stroke, wild-derived mouse strains, Neuroprotection, quantitative trait mapping

## Abstract

To identify genes involved in cerebral infarction we have employed a forward genetic approach in inbred mouse strains, using quantitative trait locus (QTL) mapping for cerebral infarct volume after middle cerebral artery occlusion. We had previously observed that infarct volume is inversely correlated with cerebral collateral vessel density in the strains. In this study, we expanded the pool of allelic variation among classical inbred mouse strains by utilizing the eight founder strains of the Collaborative Cross and found a wild-derived strain, WSB/EiJ, that breaks this general rule that collateral vessel density inversely correlates with infarct volume. WSB/EiJ and another wild-derived strain, CAST/EiJ, show the highest collateral vessel densities of any inbred strain, but infarct volume of WSB/EiJ mice is 8.7-fold larger than that of CAST/EiJ mice. QTL mapping between these strains identified four new neuroprotective loci modulating cerebral infarct volume while not affecting collateral vessel phenotypes. To identify causative variants in genes we surveyed non-synonymous coding SNPs between CAST/EiJ and WSB/EiJ and found 96 genes harboring coding SNPs predicted to be damaging and mapping within one of the four intervals. In addition, we performed RNA sequencing for brain tissue of CAST/EiJ and WSB/EiJ mice and identified 79 candidate genes mapping in one of the four intervals showing strain-specific differences in expression. The identification of the genes underlying these neuroprotective loci will provide new understanding of genetic risk factors of ischemic stroke which may provide novel targets for future therapeutic intervention of human ischemic stroke.

## Introduction

Stroke, the sudden death of the brain tissue occurring when the blood flow to the brain is lost by blockage or rupture, is a leading cause of death in the United States (the fourth) and the world (the second) (Roger *et al.* 2012; Johnson *et al.* 2016). The majority (over 80%) of strokes are ischemic in origin and results in irreversible death of brain tissue (infarction). Studies of genetic risk factors for stroke *susceptibility* in the human have employed both family-based linkage (Gretarsdottir *et al.* 2003; Helgadottir *et al.* 2004) and population-based genome-wide association studies (GWAS) (Matarin *et al.* 2007; Matarin *et al.* 2008a; Matarin *et al.* 2008b). Unfortunately, these studies have not yet uncovered druggable targets to treat stroke and modify clinical outcomes. These studies were designed to identify stroke *susceptibility* risk factors, and not factors that modulate infarct size or neurological outcomes once a stroke has occurred. More relevant to this goal would be a study of anatomic or neurological outcomes in ischemic stroke patients. A GWAS has recently been published on neurological or behavioral outcomes of 6,165 ischemic stroke patients (Soderholm *et al.* 2019) but only a single variant passed the threshold of genome-wide significance. Furthermore, despite an explosion of the human GWAS for disease phenotypes, no published GWAS for infarct volume in ischemic stroke has been published. The paucity of human genetic variants shown to modulate stroke outcomes is perhaps not surprising. Genetic studies in the human related to infarct volume or neurological outcomes are intrinsically problematic due to uncontrollable variation in the extent and location of the occluded vessel, and especially, variation in the critical time between first recognized symptoms of stroke and medical intervention. Instead, current understanding of the mechanisms underlying infarct damage is primarily based on experimental animal models.

Animal models of focal cerebral ischemia have been established to investigate the pathophysiologic events occurring after ischemic stroke. Most stroke models induce cerebral ischemia within the middle cerebral artery (MCA) territory as most relevant for thrombo-embolic stroke. MCA occlusion models vary both in the extent of occlusion (permanent *vs.* transient occlusion) and the site of occlusion (proximal *vs.* distal portion of the vessel). Several reports have determined that permanent occlusion of distal MCA method produces more restricted and reproducible damage to the cerebral hemisphere (Majid *et al.* 2000; Lambertsen *et al.* 2002; Carmichael 2005). Furthermore, previous studies have also demonstrated that different inbred mouse strains show robust differences in stroke outcomes, providing evidence that the innate response to permanent focal cerebral ischemia is under strong genetic control (Barone *et al.* 1993; Majid *et al.* 2000; Lambertsen *et al.* 2002; Sugimori *et al.* 2004).

Therefore, we have taken a forward genetic approach (quantitative trait locus – QTL – mapping) to identify novel genes/pathways (genetic components) involved in modulating infarct volume in ischemic stroke using the well-established model of permanent occlusion of distal MCA (pMCAO). Previously, we uncovered several genetic loci that are involved in regulating ischemic stroke and identified several genes modulating infarct volume within the genetic loci (Keum and Marchuk 2009; Chu *et al.* 2013; Keum *et al.* 2013; Lee *et al.* 2016; Lee *et al.* 2018).

One of these loci located on distal chromosome 7 (cerebral infarct volume QTL 1 (*Civq1*)) is the strongest and most significant locus modulating infarct volume, found in multiple pairwise crosses of inbred mouse strains (Keum and Marchuk 2009; Keum *et al.* 2013). Intriguingly, *Civq1* overlaps with a locus on chromosome 7 that modulates cerebral collateral vessel number (collateral artery number QTL 1 (*Canq1*)) (Keum and Marchuk 2009; Zhang *et al.* 2010; Keum *et al.* 2013; Lee *et al.* 2016). We and others have noted a strong inverse correlation between the volume of the infarct after pMCAO and the number of collateral vessel connections in the cerebral vasculature (Keum and Marchuk 2009; Zhang *et al.* 2010). These vessels connect portions of the same or different arteries, and upon occlusion of an artery, provide a circulatory shunt, enabling reperfusion of the ischemic cerebral territory. *Canq1* was recently shown to be due to variation in the *Rabep2* (Lucitti *et al.* 2016), encoding a protein that modulates endosomal recycling of VEGFR2, a receptor for vascular endothelial growth factor (Kofler *et al.* 2018). This locus and gene exert an overwhelming effect on the size of the infarct after vessel occlusion across a wide spectrum of inbred mouse strains.

Given the strong influence that the collateral circulation plays in the modulation infarct volume across most inbred mouse strains, we sought strains that break the inverse correlation between collateral vessel density and infarct volume. Previously, we found one such strain, C3H, which breaks this inverse correlation. Genome-wide QTL mapping for infarct volume after pMCAO in F2 progeny of a cross between B6 and C3H identified a neuroprotective locus, *Civq4*, on chromosome 8 (Chu *et al.* 2013). *Civq4* was identified in a cross between two classical inbred mouse strains that overall exhibit relatively low sequence variation compared to that found within the entire species. These and other commonly used inbred mouse strains contain 94% of their genetic background from *Mus musculus domesticus*, 5% from *Mus musculus musculus*, and less than 1% from *Mus musculus castaneus* (Yang *et al.* 2011). Thus, genetic mapping using these inbred mouse strains may systemically miss potential candidate loci that might cause phenotypic variation within the entire species. Therefore, in this study, we utilize the eight founder mouse strains of the collaborative cross (CC) mouse to expand the scope of genetic variation that we survey in the mouse genome. We use this information to select strains for QTL mapping that will uncover novel loci that modulate infarct size via a collateral-independent mechanism.

## Materials and methods

### Animals

All inbred mouse strains were obtained from the Jackson Laboratory (Bar Harbor, ME), and then bred locally to obtain mice used in all experiments. Mice (both male and female animals) were age matched (P21 for collateral vessel perfusion and 12 ± 1 week for pMCAO) for all experiments. All animal procedures were conducted under protocols approved by the Duke University IACUC in accordance with NIH guidelines.

### Collateral vessel density measurement

As collateral vessel traits are determined by 3 weeks of age and remain constant for many months (Clayton *et al.* 2008), the collateral vessel phenotype was measured at P21 as previously described (Lee *et al.* 2016; Lee *et al.* 2018). Mice were anesthetized with ketamine (100 mg/kg) and xylazine (5 mg/kg), and the ascending thoracic aorta was cannulated. The animals were perfused with freshly made buffer (1 mg/ml adenosine, 40 g/ml papaverine, and 25 mg/ml heparin in PBS) to remove the blood. The pial circulation was then exposed after removal of the dorsal calvarium and adherent dura mater. The cardiac left ventricle was cannulated and a polyurethane solution with a viscosity sufficient to minimize capillary transit (1:1 resin to 2-butanone, PU4ii, VasQtec) was slowly infused; cerebral circulation was visualized under a stereomicroscope during infusion. The brain surface was topically rinsed with 10% PBS-buffered formalin and the dye solidified for 20 min. After post-fixation with 10% PBS-buffered formalin, pial circulation was imaged. All collaterals interconnecting the anterior- and middle cerebral artery trees of both hemispheres were counted.

### Permanent MCAO

Focal cerebral ischemia was induced by direct permanent occlusion of the distal MCA as previously described (Lee *et al.* 2016; Lee *et al.* 2018). Briefly, adult mice were anesthetized with ketamine (100 mg/kg) and xylazine (5 mg/kg), and then 0.5% bupivacaine (5 mg/ml) was also administrated by injection at the incision site. The right MCA was exposed by a 0.5 cm vertical skin incision midway between the right eye and ear under a dissecting microscope. After the temporalis muscle was split, a 2-mm burr hole was made with a high-speed micro drill at the junction of the zygomatic arch and the squamous bone through the outer surface of the semi-translucent skull. The MCA was clearly visible at the level of the inferior cerebral vein. The inner layer of the skull was removed with fine forceps, and the dura was opened with a 32-gauge needle. While visualizing under an operating microscope, the right MCA was electrocauterized. The cauterized MCA segment was then transected with microscissors to verify permanent occlusion. The surgical site was closed with 6-0 sterile nylon sutures. The temperature of each mouse was maintained at 37°C with a heating pad during the surgery and then was placed in an animal recovery chamber until the animal was fully recovered from the anesthetic. Mice were then returned to their cages and allowed free access to food and water in an air-ventilated room with the ambient temperature set to 25°C.

### Infarct volume measurement

Cerebral infarct volumes were measured 24 h after surgery because the size of the cortical infarct is largest and stable at 24 h after distal permanent MCA occlusion (Lambertsen *et al.* 2005). Twenty-four hours after pMCAO surgery, the animals were euthanized, and the brains were carefully removed. The brains were placed in a brain matrix and sliced into 1 mm coronal sections after being chilled at −80°C for 4 min to slightly harden the tissue. Each brain slice was placed in 1 well of a 24-well plate and incubated for 20 min in a solution of 2% 2,3,5-triphenyltetrazolium chloride (TTC) in PBS at 37°C in the dark. The sections were then washed once with PBS and fixed with 10% PBS-buffered formalin at 4°C. Then, 24 h after fixation, the caudal face of each section was scanned using a flatbed color scanner. The scanned images were used to determine infarct volume (Wexler *et al.* 2002). Image-Pro software (Media Cybernetics) was used to calculate the infarcted area of each slice by subtracting the infarcted area of the hemisphere from the non-infarcted area of the hemisphere to minimize error introduced by edema. The total infarct volume was calculated by summing the individual slices from each animal.

### Flow cytometry

Adult (8 – 12 week) mouse brain cortex of 3 CAST and 6 WSB mice were digested with collagenase A (1.5 mg/ml) and DNase I (0.4 mg/ml) in HBSS containing 5% FBS and 10 mM HEPES at 37°C for 45 min. Pelleted post-digested brain preparations were suspended in 30% Percoll^®^ and spun at room temperature to separate cells from myelin. Isolated cells were stained with fluorophore-conjugated antibodies for 30 min at room temperature in PBS containing 3% FBS, 5 μg/ml of anti-CD16/32, 5% normal rat serum, 5% normal mouse serum and 10 mM EDTA. The antibodies were obtained from BioLegend (San Diego, CA) (anti-CD3-AF488 (145-2C11), anti-CD49b-PE (HMα2), anti-F4/80-PE-Cy7 (BM8), anti-B220-AF647 (RA3-6B2), anti-CD64-BV421 (X54-5/7.1), anti-CD103-BV510 (2E7), anti-CD45-BV605 (30-F11), anti-IA/IE-BV650 (M5/114.15.2), and anti-CD11c-BV785 (N418) or BD Biosciences (San Jose, CA) (Anti-CD31-PerCP-Cy5.5 (MEC13.3), anti-SiglecF-PE-CF594 (E50-2440), anti-Ly6G-AF700 (1A8), anti-CD11b-APC-Cy7 (M1/70), and anti-CD24-BV711 (M1/69), or Invitrogen (Carlsbad, CA) (Anti-Ly6C-PerCP-Cy5.5 (HK1.4). Dead cells were positively stained with LIVE/DEAD Fixable Yellow Dead Cell Stains (Molecular Probes, Waltham, MA). Flow cytometric analysis was performed using a LSRII flow cytometer (BD Biosciences, San Jose, CA) and results analyzed using FlowJo software (BD, Franklin Lakes, NJ).

### SNP genotyping

Genomic DNA was isolated from tails of F2 intercross between CAST/EiJ (CAST) and WSB/EiJ (WSB) mice using DNeasy Tissue kit (Qiagen, Hilden, Germany). Genome-wide SNP genotyping was performed with a Mouse Universal Genotyping Array (MUGA, 8K SNPs). Array hybridization including sample preparation was performed by Neogen/GeneSeek, Inc (Lincoln, NE).

### Linkage (QTL) analysis

Genome-wide scans were performed using R/qtl software. Genotypes from MUGA were prepared for QTL mapping as follows. A total of 2460 informative markers for CAST and WSB across the mouse genome were used for genetic mapping. The significance thresholds for LOD scores were determined by 1000 permutations using all informative markers. Each QTL was indicated as a significant when its LOD score exceeded 95% (p < 0.05) of the permutation distribution and QTL intervals exceeded 99% (p < 0.01) of the permutation distribution were considered as candidate intervals. The 95% confidence interval (CI) of each peak was determined by 1.5-LOD support interval. The physical (Mb) map positions based on the genomic sequence from the GRCm38/mm10 were calculated using Mouse Map Converter tool of the Jackson Laboratory (http://cgd.jax.org/mousemapconverter/).

### Haplotype analysis

For the intervals of the identified 4 neuroprotective loci (*Civq8* on chromosome 1, *Civq9* on chromosome 6, *Civq10* on chromosome 13, and *Civq11* on chromosome 17), SNP data were obtained from the Mouse Phenome Database (http://phenome.jax.org/). An amino acid change affected detrimental to a protein was examined by 3 independents *in silico* prediction algorithms, Polymorphism Phenotyping v2 (PolyPhen-2, http://genetics.bwh.harvard.edu/pph2/index.shtml), Sorting Intolerant From Tolerant (SIFT, http://sift.jcvi.org), and Protein Variation Effect Analyzer (PROVEAN, http://provean.jcvi.org).

### RNA sequencing analysis

Paired-end, 150 bp sequencing reads were generated from adult (8 – 12 week) brain tissue mRNA of 6 CAST (3 males and 3 females) and 6 WSB (3 males and 3 females) mice on the Illumina HiSeq 2500 platform. Adapters for all paired-end sequencing reads were trimmed using Cutadapt (Martin 2011). Differential gene expression analysis is occasionally confounded by differences in alignment rate of reads obtained from strains that are highly diverged from the reference genome. Furthermore, there are millions of variant sites between the CAST and WSB strains, which would make these effects substantially pronounced. To alleviate these effects, we performed our alignment to pseudo-reference genomes (http://csbio.unc.edu/CCstatus/index.py?run=Pseudo), produced by altering the mouse reference genome (GRCm38.p4) using variant calls for the CAST and WSB strains. Reads from each strain were aligned to their respective pseudo-reference genome using TopHat (v. 2.1.1) (Trapnell *et al.* 2009) under default settings and re-mapped to reference coordinates using the Lapels pipeline (Holt *et al.* 2013; Huang *et al.* 2014).

### Differential gene expression

Gene expression counts for each sample were obtained using HTseq (Anders *et al.* 2015) under default settings and subsequently imported into DESeq2 (Love *et al.* 2014), where gene counts for each mouse were normalized for differences in sequencing effort and dispersions were calculated for each sample. Differential expression was determined using a two-sided Wald test comparing two negative binomial distributions. *p*-values were adjusted using a false-discovery rate of 5% by Benjamini-Hochberg correction (Benjamin and Hochberg 1995).

### Statistics analysis

Results were represented as the mean ± SEM. Significant differences between datasets were determined using *p*-values *<.05* were considered significant.

### Data availability

Strain used in this study are available through the Jackson Laboratory catalog. Supplemental data contain two figures (Figure S1 and S2) and three tables (Table S1 – S3). Figure S1 shows flow cytometry data used to determine the expression level of CD45^+^ hematopoietic cells from brain cortex of CAST and WSB mice. Figure S2 shows both the number of collateral vessel connections and infarct volume after pMCAO for all individual F2 animals. Table S1 contains genotype and phenotype information of 251 F2 (CAST × WSB) animals used for QTL mapping analysis. Table S2 contains detailed information of all coding SNPs including *in silico* prediction of three independent algorithms. Table S3 contains detailed information of RNA sequencing analysis within identified neuroprotective loci. RNA sequencing data are available at GEO with the accession number: GSE129379. Supplemental material available at Figshare.

## Results

### Analysis of collateral vessel density and infarct volume after permanent MCAO for all eight founder strains of Collaborative Cross mouse strain

Using the eight founder mouse strains of the CC, we first examined the number of collateral vessel connections between the anterior cerebral artery (ACA) and middle cerebral artery (MCA) (Figure 1A and B). Classical inbred mouse strains B6, NOD/ShiLtJ (NOD), NZO/HILtJ (NZO), and 129S1/SvlmJ (129S1), and a wild-derived strain, PWK/PhJ (PWK) show similar level of vessel connections (B6 (20.4), NOD (21.0), NZO (21.4), 129S1 (22.4), and PWK (20.4)), whereas A/J (10.2) showed a lower level. Interestingly, two wild-derived strains, CAST and WSB, showed the highest number of collateral vessel connections (CAST (28.8) and WSB (27.3)) that we have ever observed in any inbred strain.

**Figure 1.**
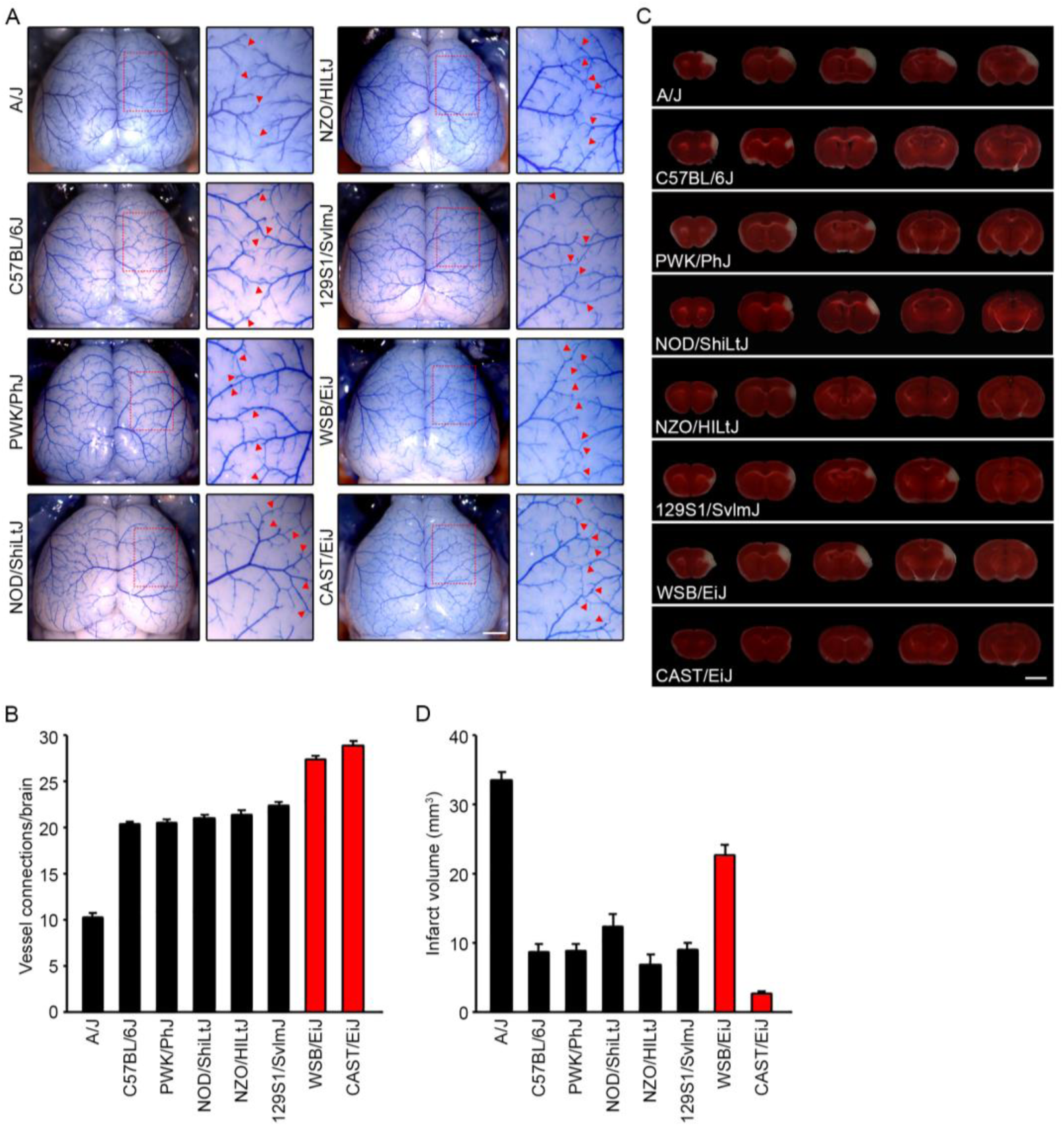
Collateral vessel density and infarct volume after pMCAO in 8 founder strains of the Collaborative Cross mouse. **(A)** Representative images of the brains for the 8 founder strains of the CC recombinant inbred mapping panel; A/J, B6, PWK, NOD, NZO, 129S1, WSB, and CAST. For each strain, the area outlined in a red box in the leftmost image is 3-fold magnified in the image to the right, where the red arrowheads indicate collateral vessel connections between the ACA and the MCA. Scale bar: 1mm. **(B)** The graph indicates the average number of collateral vessel connections between ACA and MCA in the brain. The total number of animals for; A/J, B6, PWK, NOD, NZO, 129S1, WSB, and CAST were 21, 37, 24, 9, 14, 31, 13, and 32 mice, respectively. Data represent the mean ± SEM. **(C)** Serial brain sections (1 mm) for each founder strain of the CC mouse 24 h after pMCAO. The infarct appears as white tissue after 2% TTC staining. Scale bar: 5 mm. **(D)** The graph shows the infarct volume for each founder strain of the CC mouse. The total number of animals for; A/J, B6, PWK, NOD, NZO, 129S1, WSB, and CAST were 19, 32, 27, 9, 10, 32, 18, and 32 animals, respectively. Data represent the mean ± SEM.

To determine whether collateral vessel density is inversely correlated with infarct volume in these new strains, we next examined ischemic infarct volume in the permanent occlusion model. We performed pMCAO and measured infarct volume for the eight CC founder mouse strains (Figure 1C and D). Infarct volume in these mouse strains displayed various levels after ischemic stroke induction; A/J (33.4 mm^3^), B6 (8.6 mm^3^), PWK (8.7 mm^3^), NOD (12.3 mm^3^), NZO (6.8 mm^3^), 129S1 (9.0 mm^3^), WSB (22.6 mm^3^), and CAST (2.6 mm^3^). As expected, most CC founder mouse strains showed an inverse correlation between collateral vessel connections and infarct volume. However, one strain, WSB, breaks this inverse correlation. Although two wild-derived mouse strains, CAST and WSB, exhibit the highest number of vessel connections we have seen in any inbred strain (Figure 1B), WSB consistently displayed a dramatically larger infarct volume (8.7-fold) after pMCAO compared to CAST (22.6 mm^3^ vs. 2.6 mm^3^) (Figure 1D). This suggests that at least in part, WSB modulates infarct volume after cerebral ischemia through a collateral-independent mechanism.

It is conceivable that the strain-dependent infarct volume differences are caused by different composition of cells involved in host defense mechanisms in the brains of the two strains (eg, CD45^+^ hematopoietic cells). Thus, we performed flow cytometric analysis of CD45^+^ hematopoietic cells in the cerebral cortex of the two strains. We found no difference in the fraction of microglia, lymphocytes (B and T cells), or CD11b^+^ myeloid cells (macrophages) in the brain tissue of the two strains (Figure S1 A and B).

### Four novel loci contribute infarct volume differences between CAST and WSB

In order to identify the genetic loci modulating infarct volume between these two strains, we generated F1 and F2 progeny between CAST and WSB and examined both collateral vessel density and infarct volume. Consistent with the number of collateral vessel connections between CAST and WSB, collateral vessel connections in the F1 and F2 generations exhibited a very tight distribution; WSB (27.3), CAST (28.8), F1 (27.7), and F2 (27.6) (Figure 2A). Moreover, the number of collateral vessel connections in individual F2 animals falls within a rather tight range (from 25 to 31 connections between the ACA and MCA), similar in number and range to the two parental strains (Figure S2A). Thus, genetic variation within the two strains does not appear to modulate the collateral vessel phenotype. Next, using F1 and F2 progeny, we measured infarct volume after pMCAO. In contrast to the vascular phenotype, infarct volume in the F2 generation was widely distributed, ranging from 1.1 to 38.9 mm^3^ (Figure 2B and Figure S2B) and covering nearly the entire phenotypic spectrum observed across all inbred strains (Keum and Marchuk 2009; Keum *et al.* 2013).

**Figure 2.**
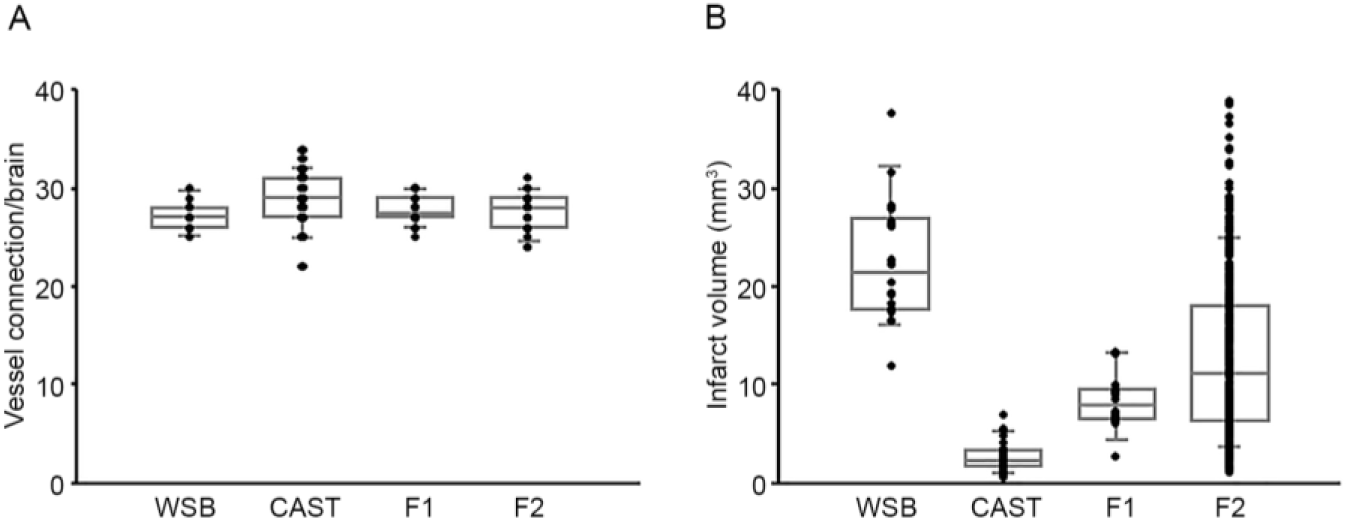
F2 intercross animals between CAST and WSB exhibit a tight distribution of collateral vessel connections but a wide distribution for pMCAO-induced infarct volume. **(A)** The dot plot graph shows the number of collateral vessel connections between the ACA and MCA for parental strains, WSB and CAST, and their F1 and F2 intercross progeny. The box plot indicates the degree of dispersion and skewness. The number of animals for WSB, CAST, F1 and F2 was 13, 32, 18, 24 animals, respectively. Data represent the mean ± SEM. **(B)** Dot and box plot graphs show the distribution of pMCAO-induced infarct volume for WSB, CAST, and their F1 and F2 intercross progeny. The number of animals for the infarct volume measurements was 18, 32, 14, and 251 animals, respectively. Data represent the mean ± SEM.

To discover genetic loci that modulate infarct volume we performed genome-wide QTL mapping analysis. A total of 251 F2 mice (Figure 2B and Figure S2B) were used for genome-wide SNP genotyping using a Mouse Universal Genotyping Array (MUGA, 8K SNPs). For genome-wide QTL mapping analysis, 2460 informative SNP markers were selected across the mouse genome (Table S1), and we identified four new QTL peaks that display highly significant linkage to the infarct volume trait. These were located on chromosomes 1, 6, 13, and 17 (Figure 3). These new loci were designated as *Civq8* through *Civq11* (Cerebral infarct volume QTL) based on our previous work that identified seven distinct loci for this trait (Keum and Marchuk 2009; Chu *et al.* 2013). The novel loci are *Civq8* (chromosome 1, LOD 5.09 (JAX00250952)), *Civq9* (chromosome 6, LOD 7.68 (UNC_rs51474193)), *Civq10* (chromosome 13, LOD 6.15 (UNC130410312)), and *Civq11* (chromosome 17, LOD 5.45 (UNC170802188)) (Table 1).

**Table 1.** Characteristics of four novel QTLs for ischemia-induced infarct volume.

| QTL | Location | LOD Score | 1.5 LOD Interval (Mb)<br>(GRCm38/mm10) | Marker at Peak<br>Peak Position (Mb) | Additive<br>Effect* | Dominance<br>Effect** | Protective<br>Allele |
| --- | --- | --- | --- | --- | --- | --- | --- |
| *Civq8* | Chr1 | 5.09 | 1: 40.66 - 68.27 (27.61 Mb) | JAX00250952<br>53.22 Mb | 6.25 | -0.77 | WSB |
| *Civq9* | Chr6 | 7.68 | 6: 22.41 - 27.88 (5.47 Mb) | UNC_rs51474193<br>26.49 Mb | -4.2 | -2.74 | CAST |
| *Civq10* | Chr13 | 6.15 | 13: 106.02 - 120.42 (14.40 Mb) | UNC130410312<br>118.60 Mb | 4.17 | -0.35 | WSB |
| *Civq11* | Chr17 | 5.45 | 17: 56.20 - 72.82 (16.62 Mb) | UNC170802188<br>61.66 Mb | -3.93 | -1.7 | CAST |

\* Direction of additive effect corresponds to additive effect of CAST allele
\*\* Direction of dominance effect corresponds to deviation from additive relationship between CAST/CAST and WSB/WSB genotypes

**Figure 3.**
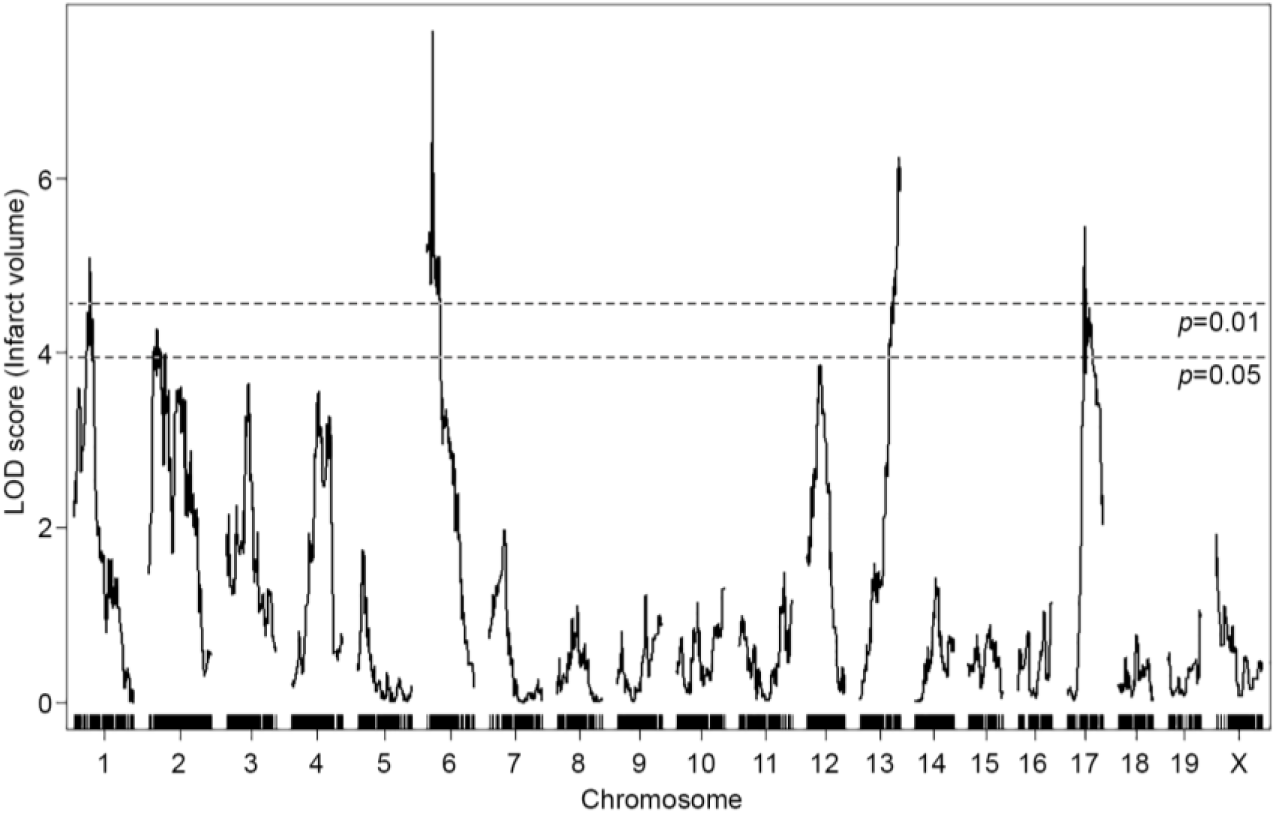
Identification of four novel *Civq* loci modulating infarct volume in the F2 intercross between CAST and WSB. The graph represents the analysis of a genome-wide QTL mapping scan for infarct volume measured 24 h after pMCAO using 251 F2 progeny (CAST and WSB). Chromosomes 1 through X are indicated numerically on the x axis. The y axis shows a logarithm of the odds (LOD) score and the significant levels (p < 0.05 and 0.01) were determined by 1000 permutation tests. Four regions of the genome mapping to chromosomes 1, 6, 13, and 17 display highly significant QTL mapping to the infarct volume trait with LOD scores of 5.09, 7.68, 6.15, and 5.45, respectively.

### New loci uncover potential candidate genes

For the precise definition of the candidate interval for each locus, we extended each interval an additional 1.5 Mb flanking each arm of 1.5-LOD support interval. One hundred seventy-three informative SNP markers were used for the definition of *Civq8* mapping to chromosome 1. The resulting candidate interval for *Civq8* is from 40.66 Mb to 68.27 Mb (Figure 4A and Table 1). Interestingly, *Civq8* harbors a transgressive allele (or alleles) that modulates the phenotype in the opposite direction of the parental strains. As shown in Figure 4B, the animals with the CAST allele are more sensitive to infarction, while those with the WSB allele are more resistant to infarction. For *Civq9*, mapping to chromosome 6, one hundred forty-four informative SNP markers were used and its candidate interval is from 22.41 Mb to 27.88 Mb (Figure 5A and Table 1). At *Civq9*, the CAST allele exhibits a protective effect on infarct volume in agreement with the phenotype of the parental CAST strain (Figure 5B). For *Civq10*, mapping to chromosome 13, one hundred twenty informative SNP markers were used and its candidate interval is from 106.02 Mb to 120.42 Mb (Figure 6A and Table 1). *Civq10* also harbors a transgressive allele (or alleles) since the WSB allele at *Civq10* is protective, in contrast to the overall phenotype of the parental strain (Figure 6B). For *Civq11*, mapping to chromosome 17, seventy-eight informative SNP markers were used and its candidate interval is from 56.20 Mb to 63.71 Mb (Figure 7A and Table 1). The CAST allele in *Civq11* exhibits a protective effect in agreement with the parental strain (Figure 7B).

**Figure 4.**
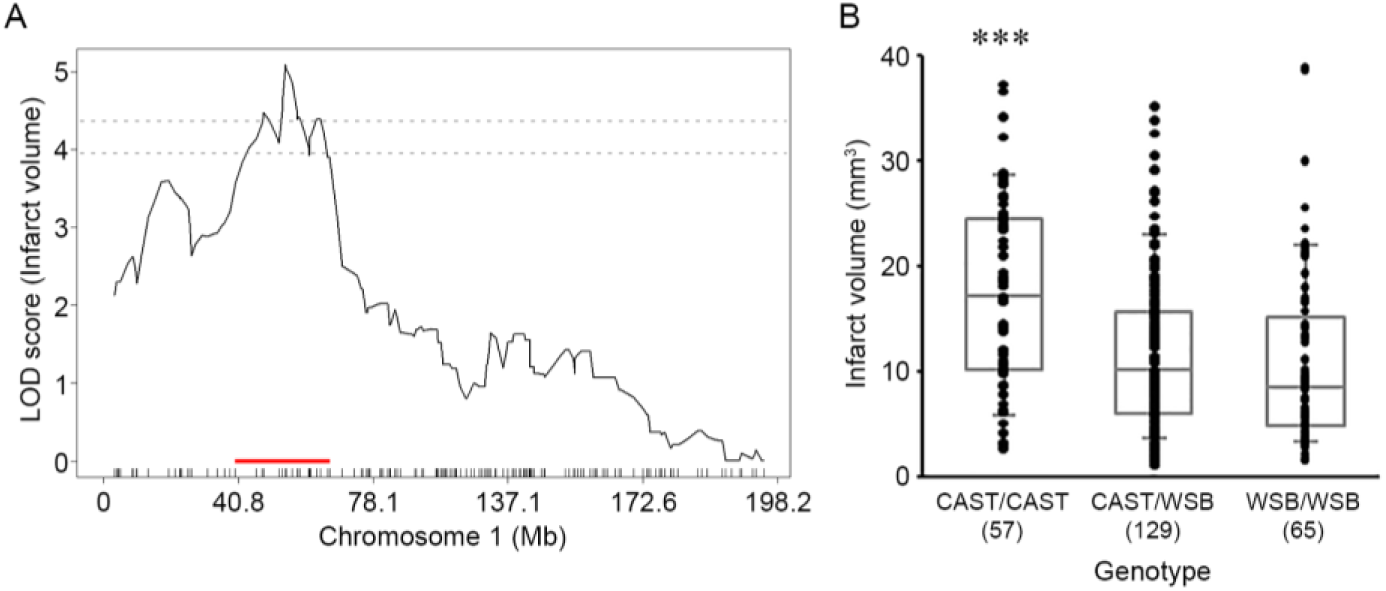
The *Civq8* locus mapping to chromosome 1. **(A)** The graph shows the QTL mapping across chromosome 1 using 173 informative SNP markers, highlighting the *Civq8* locus. The LOD score at the peak is 5.09 (JAX00250952), and the 1.5-LOD support interval is from 40.66 to 68.27 Mb, indicated by the red bar on the graph. **(B)** Genotype-phenotype correlation of the F2 cohort at JAX00250952. The alleles are transgressive with the CAST allele conferring increased susceptibility to infarction and the WSB allele conferring protection. Data represent the mean ± SEM. *** *P < .001 vs.* CAST/WSB and WSB/WSB, 1-way ANOVA followed by Scheffe’s test.

**Figure 5.**
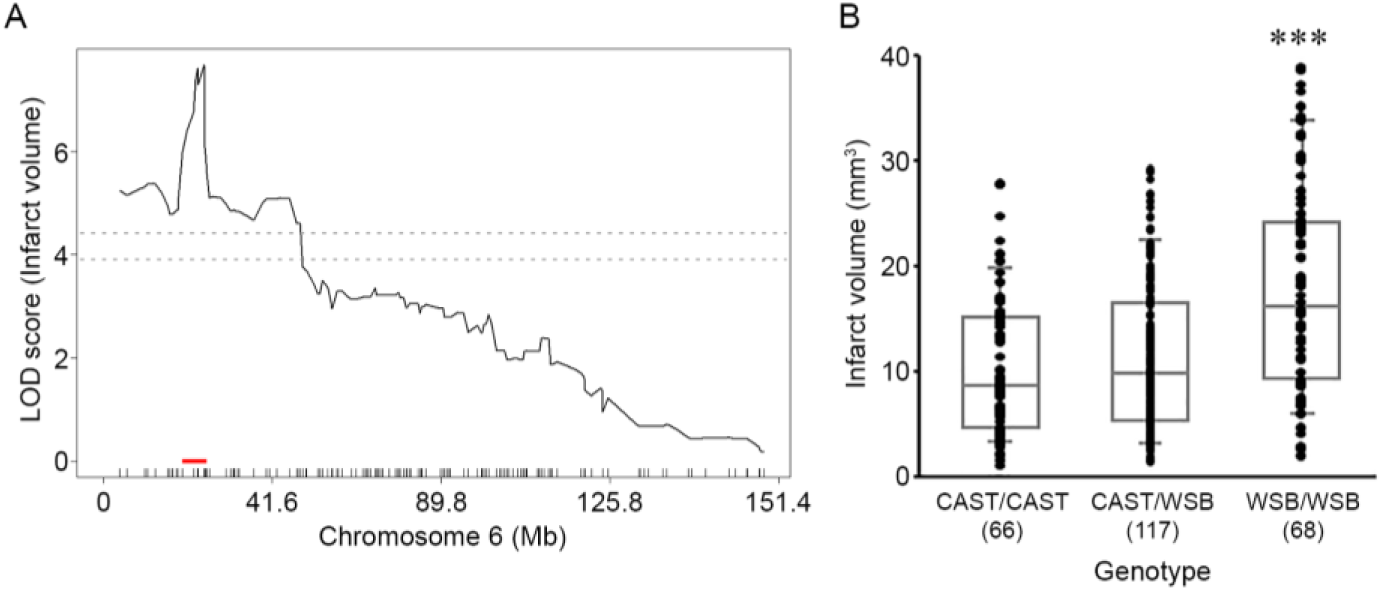
The *Civq9* locus mapping to chromosome 6. **(A)** The graph shows the QTL mapping across chromosome 6 using 144 informative SNP markers, highlighting the *Civq9* locus. The LOD score at the peak is 7.68 (UNC_rs51474193), and the 1.5-LOD support interval is from 22.41 to 27.88 Mb, indicated by the red bar on the graph. **(B)** Genotype-phenotype correlation of the F2 cohort at UNC_rs51474193. The CAST allele at *Civq9* is protective for infarction. Data represent the mean ± SEM. *** *P < .001 vs.* CAST/CAST and CAST/WSB, 1-way ANOVA followed by Scheffe’s test.

**Figure 6.**
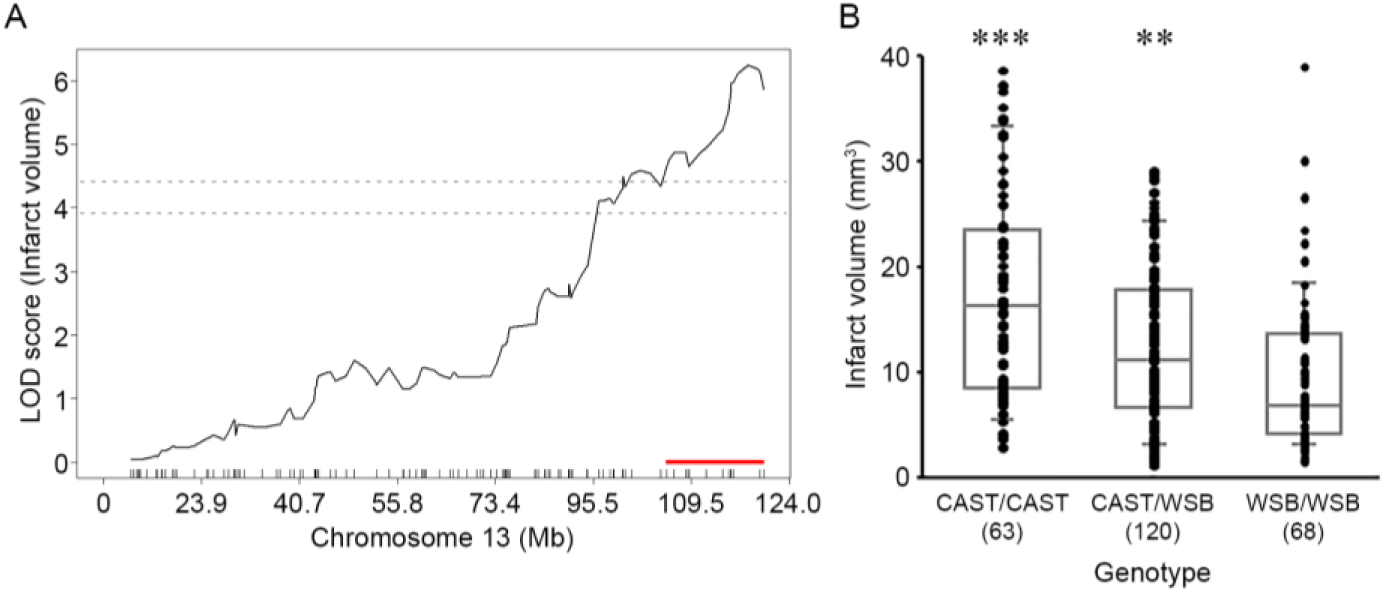
The *Civq10* locus mapping to chromosome 13. **(A)** The graph shows the QTL mapping across chromosome 13 using 120 informative SNP markers, highlighting the *Civq10* locus. The LOD score at the peak is 6.15 (UNC130410312), and the 1.5-LOD support interval is from 106.02 to 120.42 Mb, indicated by the red bar on the graph. **(B)** Genotype-phenotype correlation of the F2 cohort at UNC130410312. The alleles are transgressive with the CAST allele conferring increased susceptibility to infarction and the WSB allele conferring protection. Data represent the mean ± SEM. *** *P < .001 vs.* CAST/WSB and WSB/WSB and ** *P < .01 vs.* WSB/WSB, 1-way ANOVA followed by Scheffe’s test.

**Figure 7.**
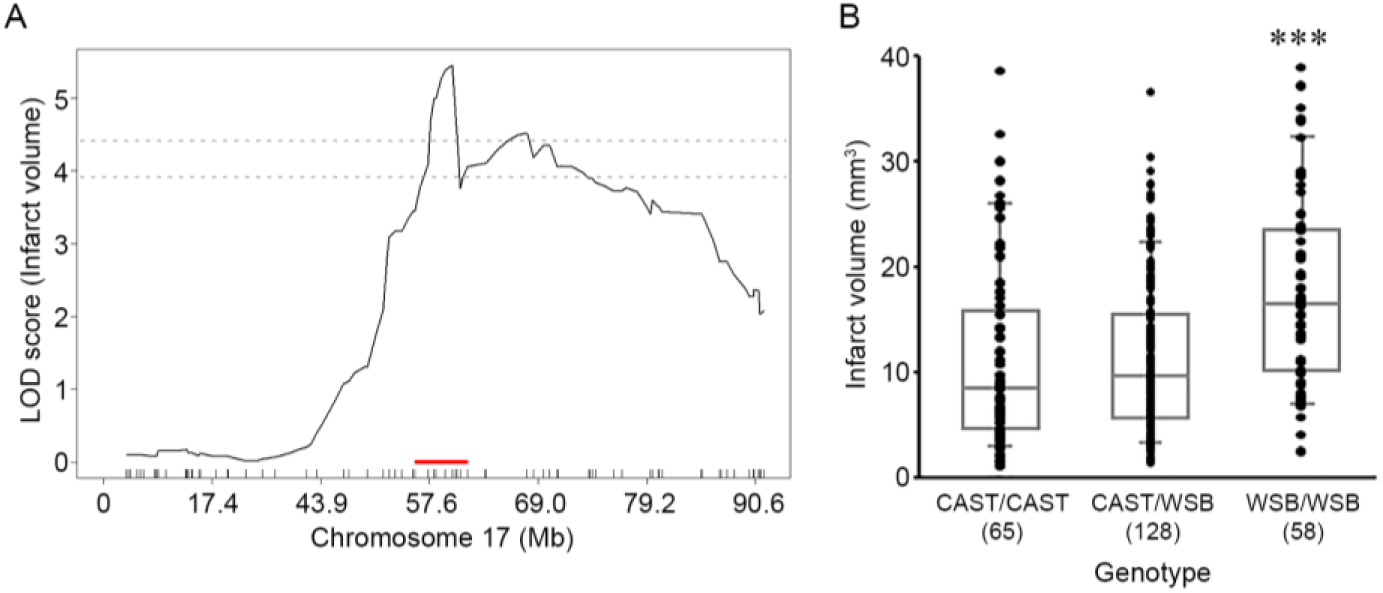
The *Civq11* locus mapping to chromosome 17. **(A)** The graph shows the QTL mapping across chromosome 17 using 78 informative SNP markers, highlighting the *Civq11* locus. The LOD score at the peak is 5.45 (UNC170802188), and the 1.5-LOD support interval is from 56.20 to 63.71 Mb, indicated by the red bar on the graph. **(B)** Genotype-phenotype correlation of the F2 cohort at UNC170802188. The CAST allele at *Civq11* is protective for infarction. Data represent the mean ± SEM. *** *P < .001 vs.* CAST/CAST and CAST/WSB, 1-way ANOVA followed by Scheffe’s test.

To identify possible candidate gene(s) modulating infarct volume in these loci, we first surveyed candidate genes within the candidate interval of each QTLs and then sought the presence of non-synonymous coding SNPs (hereafter, coding SNP) in these genes. We identified a total of 330 coding SNPs in 90 coding genes in *Civq8*, 77 coding SNPs in 20 coding genes in *Civq9*, 109 coding SNPs in 34 coding genes in *Civq10*, and 157 coding genes in 47 coding genes in *Civq11*. Within these intervals, the vast majority of these genes harbor multiple coding SNPs (Table S2). To determine whether any of these non-synonymous amino acid substitutions might affect protein function, all the coding SNPs were subjected to three independent *in silico* algorithms that predict their functional consequences; SIFT, PolyPhen-2, and PROVEAN. In *Civq8*, 46 genes have a coding SNP (or SNPs) predicted to be damaging by at least one of the prediction algorithms. However, only 11 genes harbor coding SNPs predicted to be damaging by all three programs and another 15 harbor damaging SNPs if the threshold is relaxed that only two of the three algorithms predict damaging consequences. In *Civq9*, 7 genes have coding SNPs predicted to be damaging by at least one algorithm, with only one predicted to be damaging by all three programs and another 3 genes by only two algorithms. In *Civq10*, a total 16 genes have coding SNPs predicted to be damaging with only 4 predicted to be damaging by all three programs and another 1 gene by only two algorithms. In *Civq11*, a total 27 genes have coding SNPs predicted to be damaging with only 3 predicted to be damaging by all three programs and another 7 genes by only two algorithms (Table 2 and Table S2). Although genes containing coding SNPs predicted to be damaging by all 3 independents *in silico* algorithms are the best candidate genes, even genes containing coding SNPs predicted to be damaging by only one *in silico* algorithm are still potential candidates. The details of all coding SNPs and functional predictions are listed in Table S2.

**Table 2.**
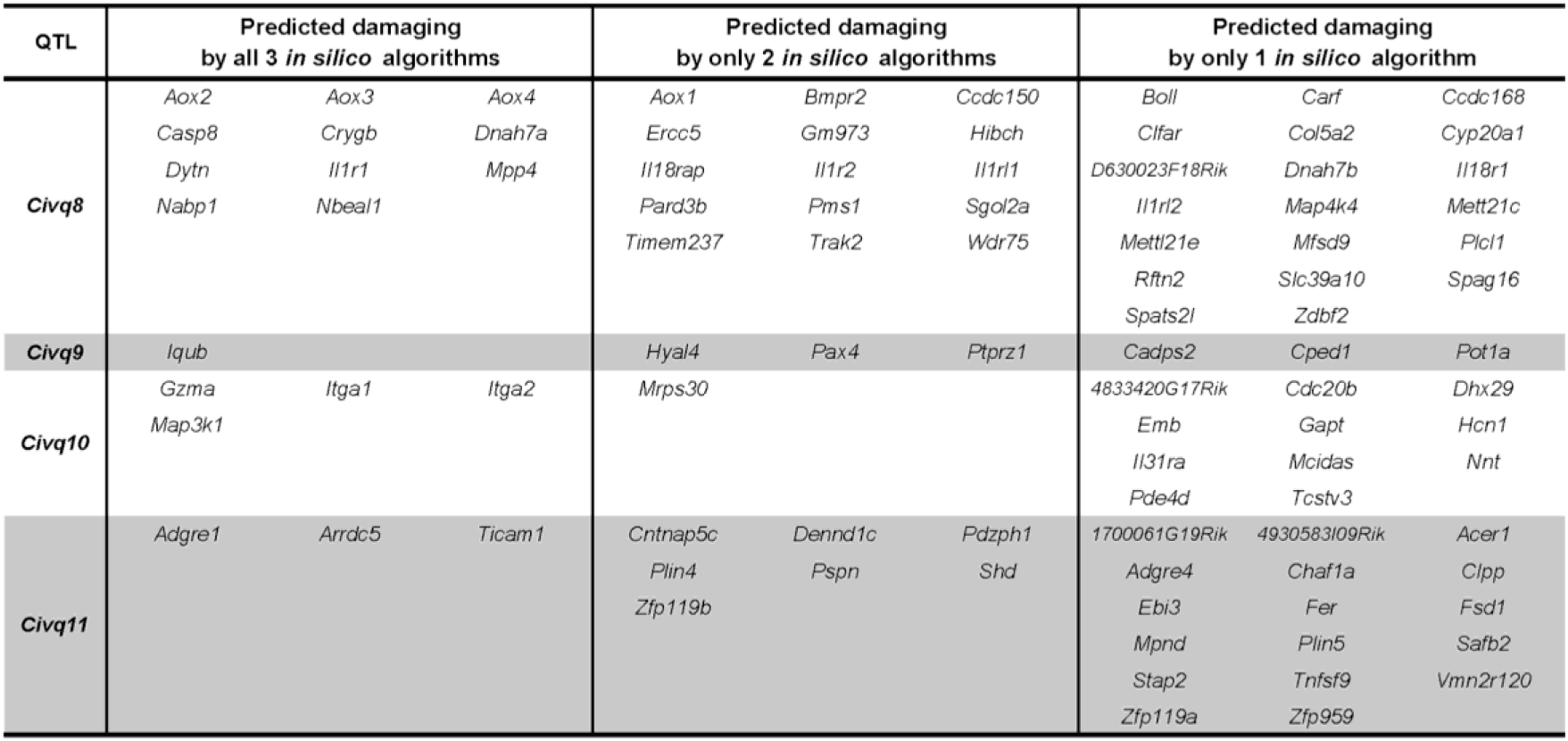
Candidate genes harboring coding SNPs that are predicted to be damaging by three different *in silico* algorithms. Detailed coding SNP information for each gene is available in Table S2.

### Strain-specific differential gene expression discovers candidate genes in the four new loci

To identify other candidate genes underlying these loci, we sought evidence of strain-specific differential transcript levels between two wild-derived mouse strains, CAST and WSB. Strain-specific differences in transcript levels could potentially be caused by regulatory sequence variation acting in *cis* by any number of potential molecular mechanisms. We employed RNA sequencing data analysis using adult brain tissues between CAST and WSB. As shown in Figure 8, a total 9,563 genes were differentially expressed between strains but only 220 of these genes are located within one of the four new QLTs, with 135 in *Civq8*, 10 in *Civq9*, 47 in *Civq10* and 28 in *Civq11*. Assuming that the most likely candidate genes would show at least a 2-fold difference in strain-specific expression reduces the overall list to only 79 candidate genes; 47 genes in *Civq8*, 3 genes in *Civq9*, 19 genes in *Civq10*, and 10 genes in *Civq11* (Table 3A and B). All 220 genes are displayed in Table S3 which includes information on direction of difference, that is, which inbred strain of the pair exhibits the higher expression. Genes harboring either coding SNPs predicted to be damaging or those showing differential expression between CAST and WSB are potential candidate genes that modulate infarct volume after ischemic stroke via a collateral vessel-independent mechanism.

**Table 3.**
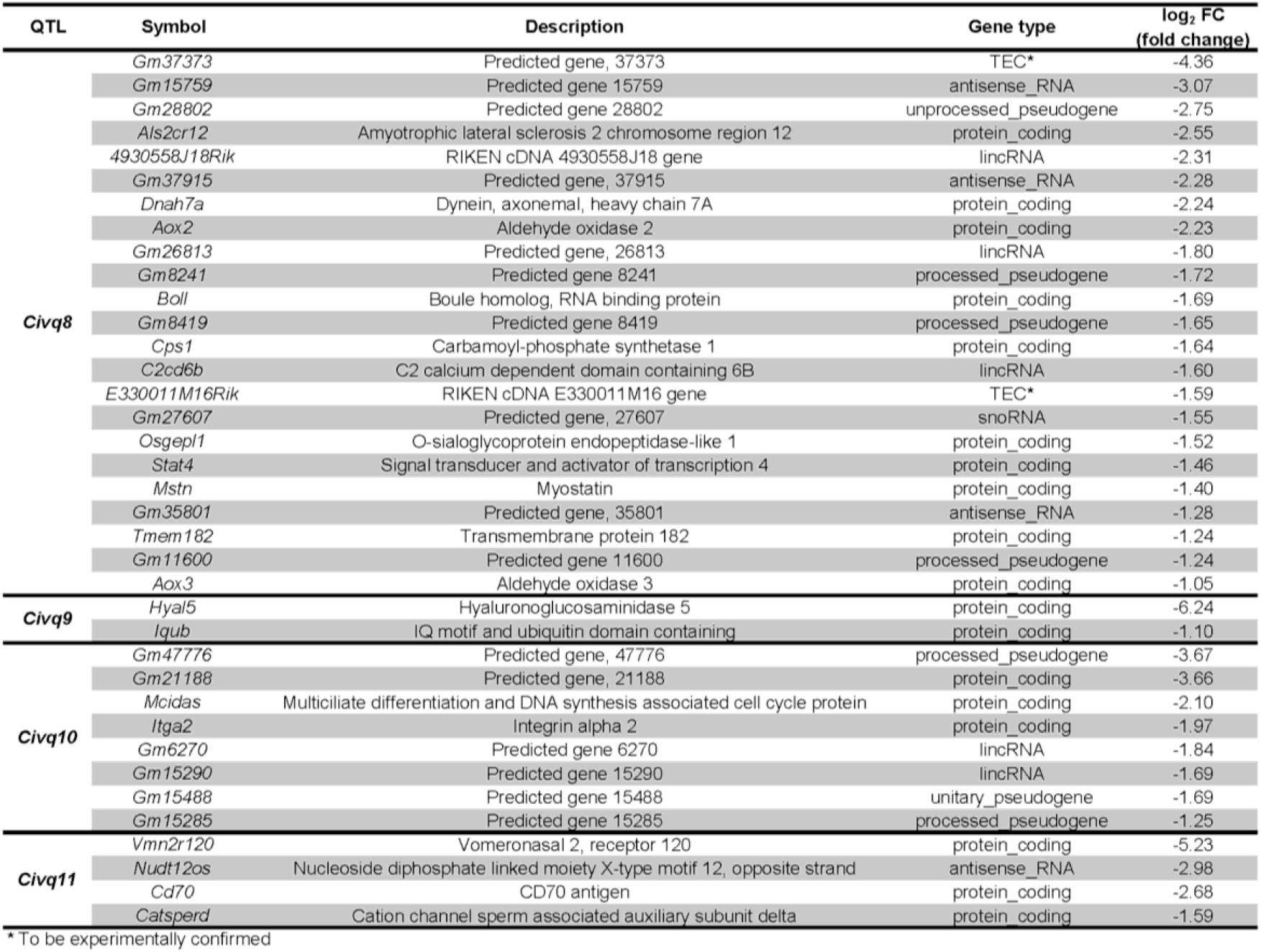
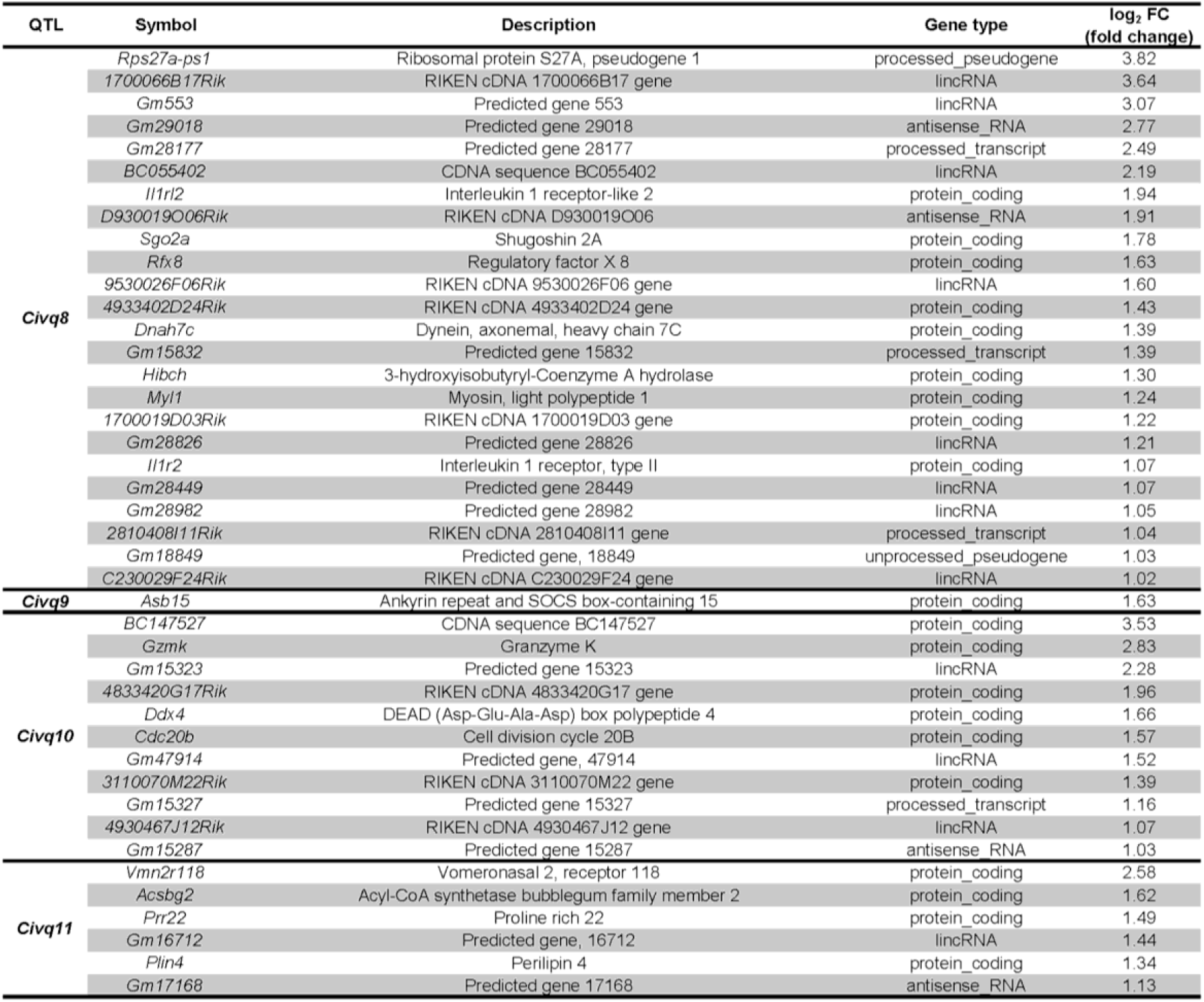
Differential gene expression between CAST and WSB determined by RNA sequencing analysis. **(A)** The table lists all genes showing at least 2-fold differential expression where the CAST allele is higher than WSB allele. **(B)** The table lists all genes showing at least 2-fold differential expression where the WSB allele is higher than CAST allele. Additional information for all 220 genes is available in Table S3.

**Figure 8.**
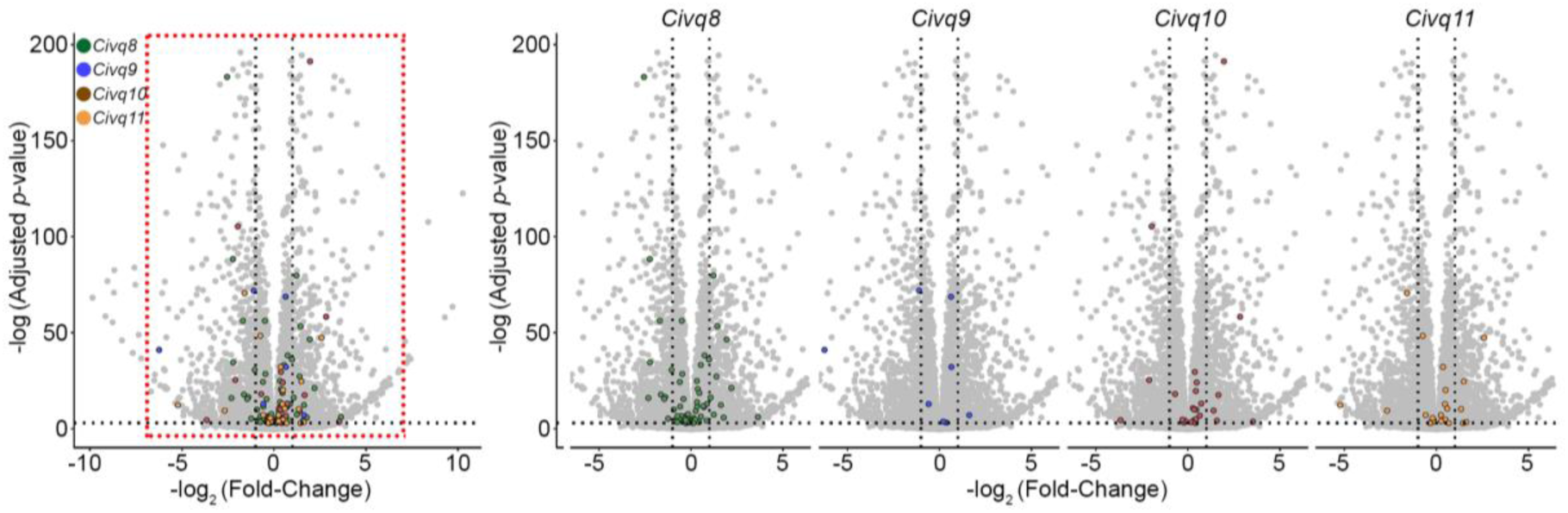
RNA sequencing data identifies genes showing strain-specific differences in cerebral cortex gene expression. **(A)** The volcano plot shows differential gene expression between CAST and WSB from brain cortex determined by RNA sequencing analysis. Each dot represents a different gene with the log2 fold change plotted against log10 *p*-value. The total number of significantly differential expressed genes across the mouse genome is 9,563 genes but only 220 genes map within one of the four QTL intervals. Differentially-expressed genes in the red box in the leftmost plot are separated for each interval on the right; green dots indicate 135 genes mapping within *Civq8*, blue dots indicate 10 genes mapping within *Civq9*, brown dots indicate 47 genes mapping within *Civq10*, and orange dots indicate 28 genes mapping within *Civq11*.

## Discussion

The long-term objective of this work is to identify novel targets for stroke therapy, focusing on those factors that modulate the size of the cerebral infarct in a collateral vessel-independent manner. In support of this goal, a recent study of a meta-analysis of multiple clinical stroke cohorts showed that collateral vessel anatomy did not correlate with neurological outcomes (de Havenon *et al.* 2019). Furthermore, the collateral vasculature is developmentally established and consequently, may be difficult to modify therapeutically.

Here, to increase the genetic diversity of our mapping approach, we utilized eight founder mouse strains of the CC recombinant inbred mapping panel. These strains include five classical inbred strains (A/J, B6, NOD, NZO, and 129S1) and three wild-derived strains from Mus musculus subspecies (PWK, WSB, and CAST). Specifically, these wild-derived strains, PWK (*musculus*), WSB (*domesticus*), and CAST (*castaneous*), are more genetically diverse than the classical inbred strains (Churchill *et al.* 2004; Threadgill 2004; Aylor *et al.* 2011; Collaborative Cross Consortium 2012). Using all eight founder strains of the CC mouse, we surveyed both the number of collateral vessel connections as well as infarct volume after pMCAO. The number of collateral vessel connection of two wild-derived strains, CAST and WSB, were the highest we have observed in any inbred mouse strains. Consistent with the role of collateral vessels in reperfusion of the ischemic territory following pMCAO, CAST mice exhibit the lowest infarct volume that we have ever measured. However, although CAST and WSB have the largest number of collateral vessel connections, unlike CAST, upon pMCAO, WSB exhibits a large infarct volume. Therefore, this wild-derived WSB strain also breaks the inverse correlation between collateral vessel connections and infarct volume. Supportive evidence for a non-collateral mechanism is the observation that infarct volume in the F2 animals shows a broad range of values while the collateral vessel connections are nearly invariant among F2 animals. Based on the ability to reperfuse the ischemic tissue due to more than sufficient collateral vessel connections, but displaying a much larger infarct volume than would be predicted, WSB likely contains loci that upon ischemic insult, contribute to neuronal death. One compelling hypothesis is that most strains contain natural neuroprotective loci that are defective in WSB. In this F2 cross, we identified the most significant four novel neuroprotective loci (*Civq8* on chromosome 1, *Civq9* on chromosome 6, *Civq10* on chromosome 13, and *Civq11* on chromosome 17). Importantly, these loci identified in an F2 cross between CAST and WSB did not overlap with other loci that we previously identified with other F2 crosses. More importantly, the *Civq1* / *Canq1* locus, the strongest locus controlling collateral vessel number, is absent in this cross. Interestingly but not surprisingly, we found transgressive alleles among the loci, alleles that operate in the opposite direction of the parental strain phenotypes. Transgressive segregation is not uncommon in intraspecific crosses (Palijan *et al.* 2003; Roper *et al.* 2003), and one of our other, previously-mapped neuroprotective loci, *Civq4*, also harbors a transgressive allele (Chu *et al.* 2013).

From the QTL mapping analysis for infarct volume, we identified four candidate intervals regulating infarct volume through a collateral-independent mechanism. In order to identify candidate genes within the 1.5 LOD-support intervals, we searched for coding SNP differences between CAST and WSB focusing on genes where the functional consequence of the variants were predicted using three different *in silico* prediction algorithms (Ng and Henikoff 2003; Adzhubei *et al.* 2010; Choi *et al.* 2012). Coding SNPs predicted as “damaging” by all 3 algorithms were considered the highest priority candidates, with full realization that any of the other genes harboring coding SNPs cannot be ignored.

As the genes underlying these four loci might harbor *cis*-regulatory sequence variation modulating differences in mRNA levels, we determined strain-specific differential transcript levels between CAST and WSB using RNA sequencing analysis. Genes exhibiting a statistically significant, 2-fold or greater difference in transcript levels are considered our highest priority candidate genes.

Through these two independent approaches, we have narrowed our list of candidate genes for all four novel loci to ninety-six genes harboring potentially damaging coding SNP variation and seventy-nine genes exhibiting greater than two-fold, strain-specific differential gene expression between CAST and WSB. To identify the causative genes, further functional studies are required. However, with additional bioinformatics analyses and a search of the relevant literature, two genes stand out.

*Civq8* harbors a protective WSB allele. *Casp8* (caspase 8), mapping within the interval, harbors multiple coding SNPs for which the WSB allele of one SNP in particular, (rs226995171), is predicted to be damaged by all three algorithms used in our analysis. Moreover, among 37 inbred mouse strains (Sanger4: Sanger SNP and indel data, 89+ million locations, 37 inbred strains of mice (2017)), only the CAST and SPRET/EiJ strains harbor this rare allele. *Casp8* initiates an extrinsic pathway during apoptotic cell death (Kischkel *et al.* 1995; Thornberry and Lazebnik, 1998) and it is possible that a poorly functioning or non-functional *Casp8* would protect brain tissue against ischemia-induced apoptotic cell death.

The other gene is *Map3k1*, a member of the mitogen-activated protein kinase kinase kinase (MAP3K) superfamily controlling the MAPKK-MAPK signaling cascade. Although MAP3K1 plays an important role in multiple aspects of cell physiology, MAP3K1 also regulates apoptosis in response to multiple cellular stresses (Xia *et al.* 1995; Cardone *et al.* 1997; Deak *et al.* 1998; Widmann *et al.* 1998). Similar to *Civq8, Civq10* is also a transgressive locus where the WSB allele is protective. One of the coding SNPs (rs257092960) in *Map3k1* is predicted to be damaged by all three algorithms and WSB is the only strain harboring this allele among 37 mouse inbred strain (Sanger4). As with *Casp8*, it is possible that a non-functional *Map3k1* protects ischemia-induced apoptotic cell death.

In summary, we have identified a wild-derived mouse strain, WSB, which breaks the inverse correlation between collateral vessel connections and infarct volume after pMCAO. By selecting two wild-derived mouse strains, CAST and WSB, that contain similarly high numbers of collateral vessel connections but which display large differences in infarct volume, we identified a strain pair that might uncover novel loci involved in neuroprotection. In a cross between these strains, we discovered four novel neuroprotective genetic loci. Using RNA sequencing of brain tissue and *in silico* analyses of coding SNPs, we further prioritized the genes mapping within these intervals. The identification of such neuroprotective genes may provide novel targets for future therapeutic intervention for human ischemic stroke.

## Acknowledgments

The authors thank Dr. Sena Bae and Mr. Daniel A. Snellings for helpful discussion concerning data sorting and analysis. This work was supported by a grant from NIH grant 5R01HL097281 and the Foundation Leducq Transatlantic Network of Excellence in Neurovascular Disease (17 CVD 03).

